# Combining AlphaFold 3 with microscale thermophoresis for aptamer optimisation

**DOI:** 10.64898/2026.10.08.757587

**Authors:** Ákos Harkai, Tamás Mészáros

**Affiliations:** Department of Molecular Biology, Institute of Biochemistry and Molecular Biology, Semmelweis University, Tűzoltó street 37-47., 1094 Budapest, Hungary

**Keywords:** aptamer, AlphaFold 3, biofilm, glucan binding protein C, interaction, microscale thermophoresis, prediction

## Abstract

Although use of AI-based tools has gained ground in molecular biology research, their reliability remains a matter of debate. Here, we combined AlphaFold 3 modelling with microscale thermophoresis to investigate the interactions between glucan-binding protein C and the biofilm-inhibiting aptamer and its five derivatives. Structural modelling identified a binding motif on the aptamer and suggested a mechanism of action; however, the predictions were only partially supported by the experimental results. The AlphaFold generated confidence metrics did not correlate directly with the measured binding affinities, indicating that confidence scores can help identify plausible interaction interfaces, but they are not reliable predictors of binding strength. These results highlight the importance of integrating structural prediction with biophysical validation to dissect aptamer–protein recognition.

**Graphical abstract:** Graphical abstract presents the workflow of our study from the initial to the further steps. Plus (+) symbolizes the ‘and’ relationship between the prediction and measurement.

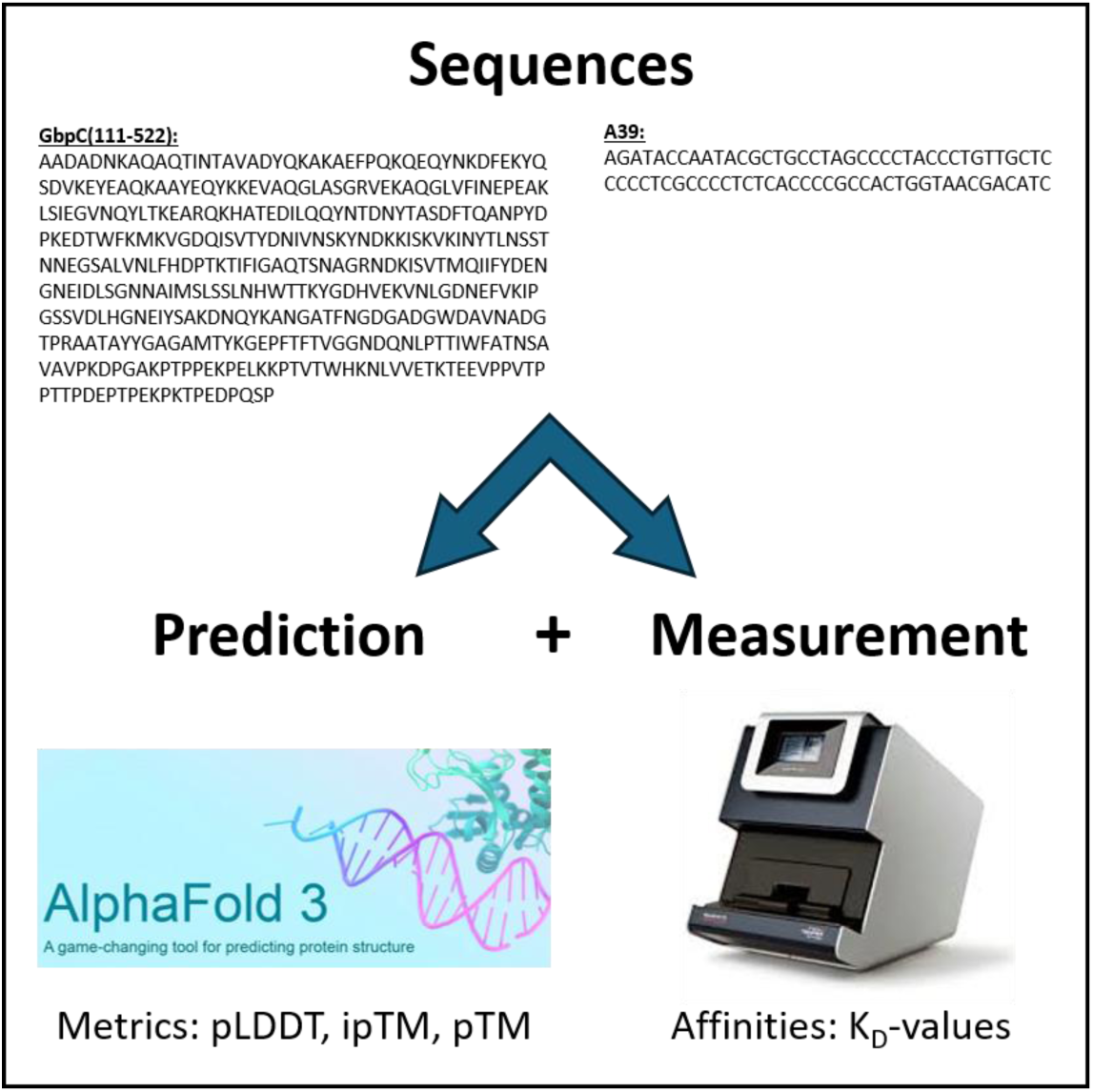

## 1. Introduction

Aptamers were first described as synthetic oligonucleotides that bind their target molecules with remarkable affinity and specificity [1,2]. As their selection methods and structural properties were developed, their wide-ranging practical significance soon became apparent [3]. Aptamers may function as bioreceptors used in diagnostics or as components of routine reagents [4–7]. Besides these applications, aptamers developed for therapeutic purposes also show great promise as drug candidates [8–10].

The most common structural biology methods for studying aptamer-target interactions are X-ray crystallography, NMR spectroscopy and cryo-EM. Crystallography enables high-resolution data of complexes, if the interacting members are available in high purity and form a well-diffracting crystal. Aptamer folding can be monitored in solution using NMR, but the upper limit on molecular weights and the insoluble partner may compromise the reliability of obtained results. In contrast, cryo-EM can provide high-resolution structural information even for large complexes without requiring the challenging crystallisation process, though it relies on dedicated laboratories with advanced instrumentation and computational capabilities [11]. Among the thousands of published aptamers collected in databases [12,13], only a few dozen possess experimentally determined 3D structure [14]. In most cases, the complex formation of newly selected aptamers and targets is barely explored, even though deciphering their spatial structures could facilitate the identification of aptamers intended for specific applications [15]. This gap raises the scientific need for a cost-effective, user-friendly and easy-to-access approach to the analysis of aptamer-target complexes.

The launch of AlphaFold has revolutionised the field of bioinformatic modelling of molecular structures in recent years. The platform employs machine learning algorithms and neural network architectures trained on experimentally determined molecular structures. By incorporating physical and biological principles driving protein folding and stability, it enables highly-accurate prediction of three-dimensional protein confirmation [16–18]. In AlphaFold 3 (AF3), the capabilities of AlphaFold have been expanded to include ions, post-translational modifications into prediction, and to model biomolecular interactions, such as protein-ligand, protein-protein and protein-nucleic acid interactions [19]. Although the impact of AF3 on structural biology is beyond doubt, the extent to which its predictions reflect the actual binding affinity of interacting partners remains unclear, especially in the context of aptamer-target interactions.

To validate AF3-predicted aptamer–target interactions against experimentally determined kinetic parameters, it is preferable to employ a technique that operates in a homogeneous solution and does not require immobilization of either binding partner. Such approaches yield equilibrium dissociation constants that more closely reflect physiological conditions. Microscale thermophoresis (MST) meets this criterium and has proven suited for investigating a wide range of biomolecular interactions, including those involving aptamers and proteins [20–23]. The principle of MST is based on the fact that binding events alter the physicochemical properties of the interacting molecules, including their total mass, charge and solvation shell [24]. These changes affect the thermophoretic behaviour of the molecules in a temperature gradient, which can be detected as variations in fluorescence signals.

One of our recently selected aptamers targeting streptococcal Glucan-binding protein C (GbpC) has proved effective in inhibiting bacterial biofilm formation [25]; however, the binding site of the lead aptamer and the structure of their complex remain unknown. In this study, we compare the confidence metrics generated by AlphaFold 3 for aptamer–GbpC complex models with binding affinities determined by microscale thermophoresis to evaluate the relationship between computational predictions and experimental measurements, and to identify potential structural and sequence determinants of the A39 aptamer–GbpC interaction.

## 2. Materials and methods

### 2.1. Aptamer derivatives

We used the chemically synthesised GbpC-specific A39 lead aptamer published previously [25] and its five created derivatives for both in silico and in vitro binding assays. The oligonucleotides and their properties are listed in Table 1. The guanine-cytosine content (GC%) and the melting temperature (T_M_) of oligonucleotides were determined by Eurofins’ Oligo Analysis Tool [26].

**Table 1.**
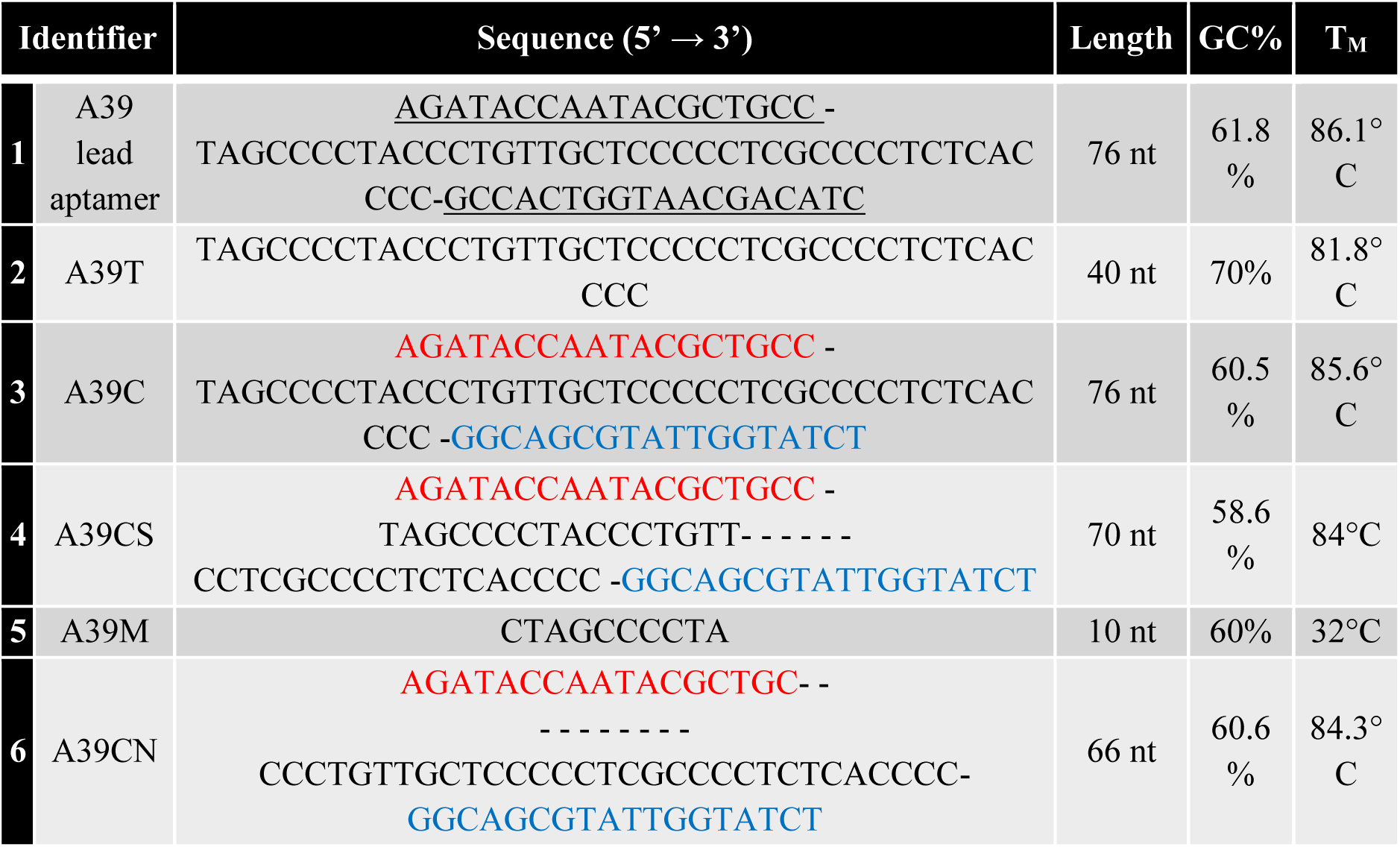
Oligonucleotides and their properties used for binding assays.

### 2.2. In silico binding assay with AlphaFold 3

Each DNA oligonucleotide was analysed separately with truncated GbpC (aa: 111-522) target protein derived from *Streptotoccus mutans* (UniProt: <u>Q8DTF1</u>) using AlphaFold 3 Server. To ensure the native-like milieu of complex formations Na^+^, K^+^, Mg^2+^, Ca^2+^ and Cl^−^ were added to sequences as input for predictions. The spatial structures of predicted complexes were assessed according to their binding sites and confidence metrics such as pLDDT, ipTM, pTM [19]. Figures were rendered using the PyMOL Molecular Graphics System (v3.1, Schrödinger, LLC.).

### 2.3. Overexpression and purification of GbpC

The GbpC target protein was produced and purified as described previously [25]. Briefly, the N-terminal 6xHis-tagging pET-28a-GbpC^(111-522)^ construct was transformed into *E. coli* BL21(DE3) competent cells (New England Biolabs) in accordance with the manufacturer’s instructions [27]. The overexpression of GbpC was conducted by auto-inducing protein production approach [28]. To this end, positive colonies were added to MDG non-inducing medium supplemented with 50 µg/ml kanamycin (Gibco). The cell culture was grown in sterile tube (Sarstedt) at 37°C for 5 h with 220 rpm rotation in an aerobic incubator. Then, 50 µl of non-induced culture was inoculated into 100 ml ZYM-5052 auto-inducing medium supplemented with 50 µg/ml kanamycin (Gibco). The induced culture was grown aerobically in sterile glass flask at 37°C for 16 h with 220 rpm rotation. Afterwards, the auto-induced medium was split into five sterile conical tubes and centrifuged at 4°C for 20 min at speed of 4000 rpm. Supernatants were then discarded and centrifuged bacterial pellets were stored in a −20°C freezer until the purification process.

Overexpressed recombinant GbpC was purified using PureCube INDIGO Ni-MagBeads (Cube Biotech) following the manufacture’s protocol [29]. In order to avoid protein degradation during isolation, protease inhibitor cocktail (Sigma, P8849) was added to the binding buffer. Purified protein was then eluted from magnetic beads and buffer-exchanged to 1x Artificial Saliva Buffer (ASB) [25] using 30 kDa MWCO Amicon ultra centrifugal filter (Merck) for in vitro binding assays. The purity of protein fractions was verified by SDS-PAGE and quantified using NanoDrop 1000 (Thermo) spectrophotomer at 280 nm.

### 2.4. In vitro binding assay with MST

Interaction measurements were performed using microscale thermophoresis (MST). A39 derivatives were synthesized with 5’-Cy5 fluorescent labelling (Eurofins). Buffer-exchanged 200 µM GbpC stock solution was diluted in a 16-step serial dilution, 1:1 (V/V) with 1xASB. Each dilution step was added with an equal volume of 100 nM 5’-Cy5-labelled oligonucleotide in 1xASB + 0.02% Tween-20 (Sigma) solution. After 5 min incubation at room temperature, reaction mixtures were loaded into Monolith Capillaries (NanoTemper). Red LED power was set to 20% for excitation, while the MST power was set to 40%. The fluorescence signal was followed before, during and after infrared laser had been turned on for 3, 20 and 1 sec, respectively using the Monolith NT.115 (NanoTemper) instrument at 25°C. Obtained MST traces were evaluated and dissociation constants were determined using MO.Affinity Analysis software (v2.3, NanoTemper). Fraction bounds were rendered with GraphPad Prism 10 (v10.3, GraphPad Software, LLC.).

## 3. Results

### 3.1. Evaluation of predicted complex structures

To determine the spatial structure of the previously selected aptamer A39 in complex with the GbpC protein, we performed in silico structural predictions using AlphaFold 3 (Fig. 1A). The resulting model suggested that neither primer region, apart from a single nucleotide from the 5′ primer, nor a substantial portion of the random region participates in interactions with the protein partner. Considering these findings, we designed five variants of the original aptamer. These included a primer-truncated version (A39T), only the nucleotides, which are involved in binding of A39 (A39M), and a construct containing an altered, fully complementary primer sequences (A39C) to promote a more stable intrinsic three-dimensional aptamer structure. In addition, two further derivatives of A39C were generated: one containing a shortened random region (A39CS), and another lacking the nucleotide motif identified as directly involved in GbpC binding in the original A39C aptamer (A39CN).

**Figure 1.**
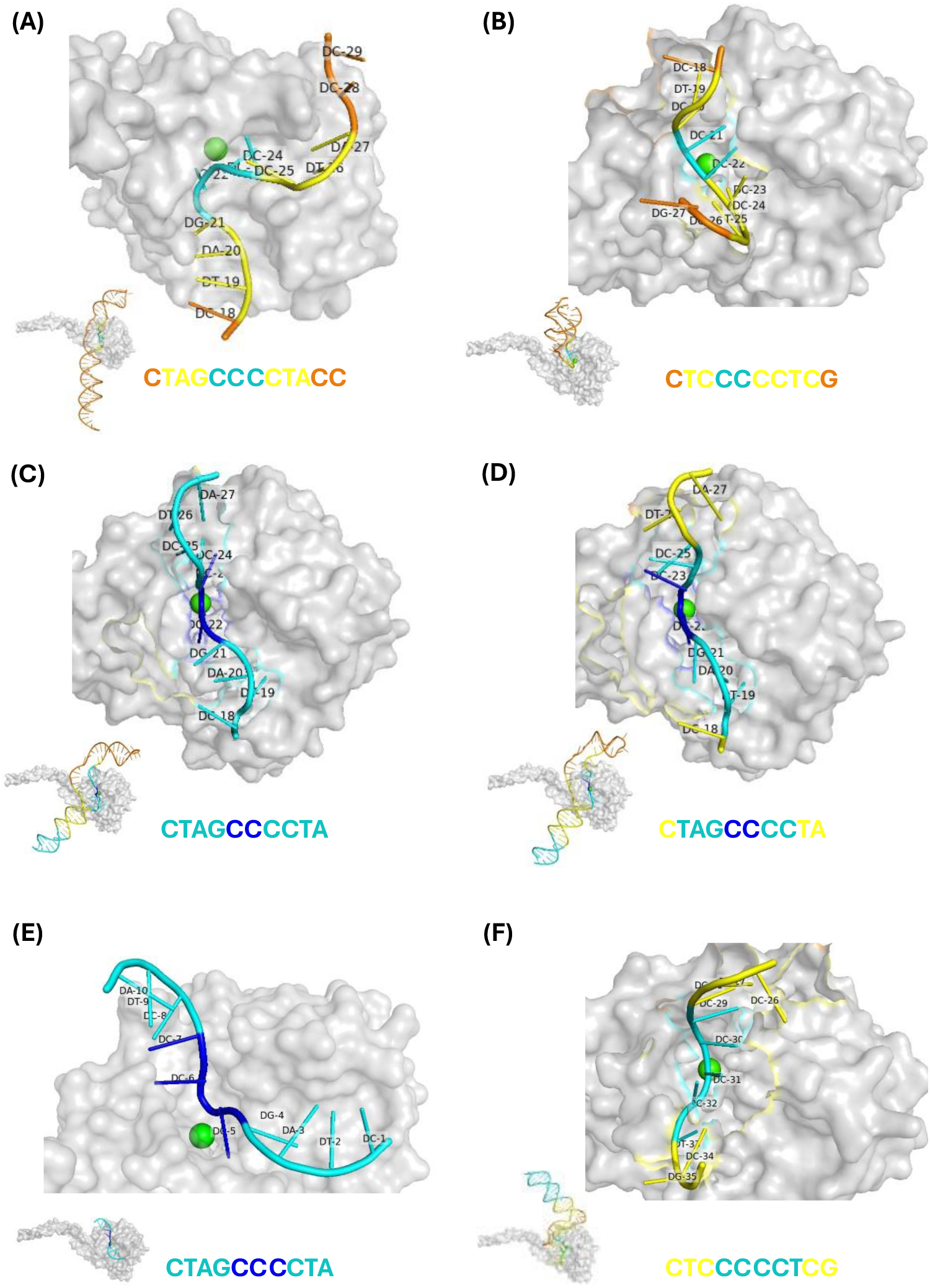
Model structures of GbpC-aptamer’s derivatives predicted by AlphaFold 3. Each derivative was modelled in complex with calcium ion (green) cofactor-binding GbpC (grey) and the overall prediction was demonstrated in left down. The protein-nucleic acid interfaces are shown enlarged and binding sites are highlighted in accordance with the pLDDT colour-coding. GbpC was modelled with; A39 lead aptamer (A), truncated A39T (B), complementary primer-holding A39C (C), complementary primer-holding with a shortened loop region A39CS (D), only motif A39M (E) and complementary primer-holding without motif A39CN (F), respectively.

AF3 analysis of the six aptamer-protein complexes showed notable differences in model confidence and conformational stability. The original A39 aptamer and its primer-truncated variant (A39T) exhibited relatively low per-residue confidence scores, indicating the presence of flexible and poorly defined regions within the predicted structures (Fig. 1A, B, and Table 2). In contrast, the complementary primer region holding derivatives (Fig. 1C, D, F) displayed higher local structural confidence, as reflected by their pLDDT score distributions (Table 2). All variants containing complementary primer sequences adopted a stable hairpin-like architecture with a single unstructured loop region (Fig. 1C, D, F). Collectively, these predictions suggest that the aptamer derivatives recognize and bind the target protein through distinct structural conformations.

**Table 2.**
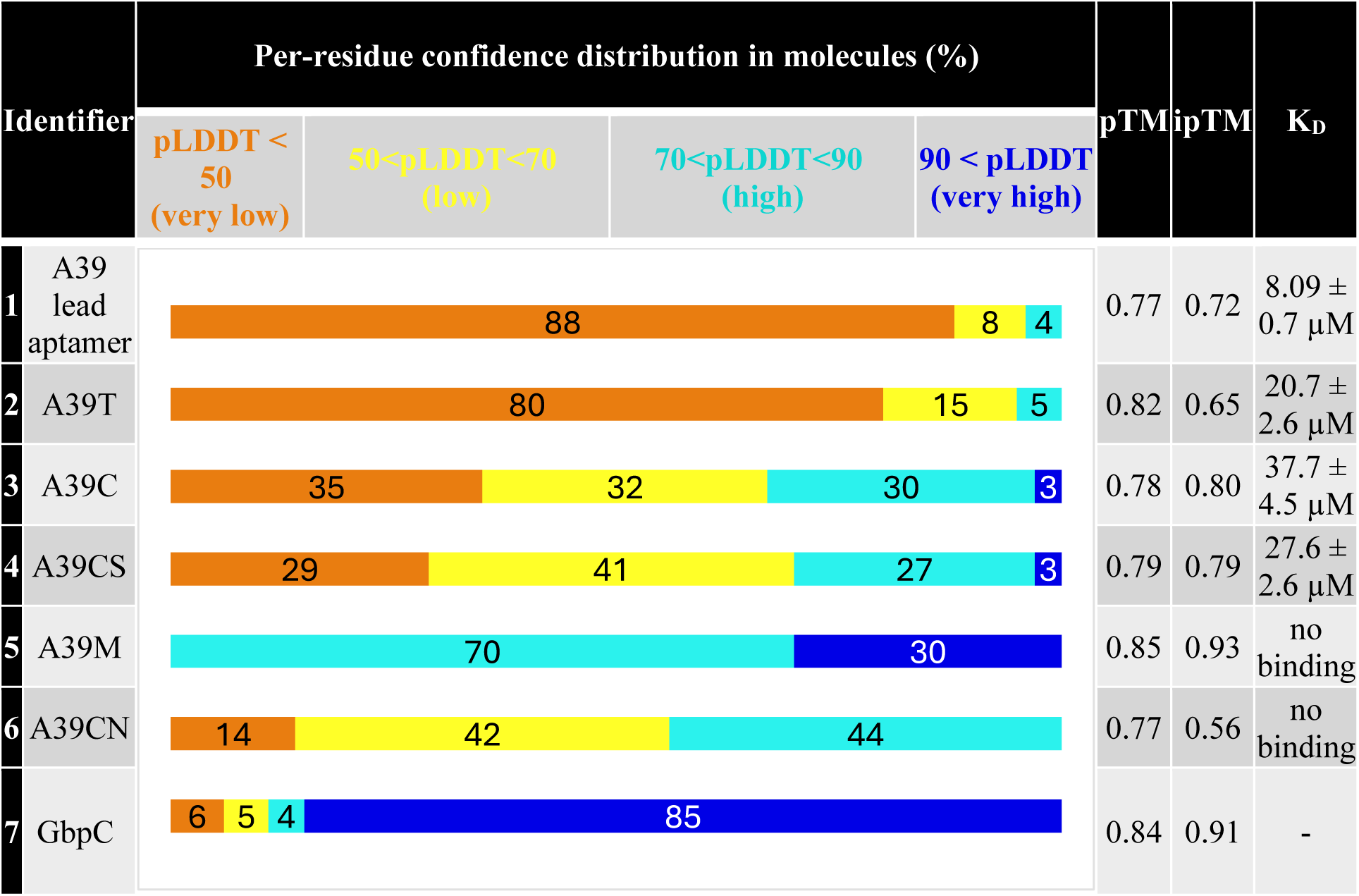
Summary of the confidence metrics predicted by AlphaFold 3 and the affinity results of MST measurements.

Next, we aimed to investigate the amino acid and nucleotide motifs of GbpC and the aptamers, respectively, that are directly involved in mediating the interactions. To this end, we screened the predicted models for oligonucleotide segments with the highest pLDDT values, that refers to confident backbone and side-chain orientations. Without exception, all identified 10-12 nucleotide-long segments formed binding interfaces within an internal groove of GbpC in the vicinity of calcium ion (Fig. 1), despite of the presence of a more exposed electropositive protein surface that might be expected to favour aptamer recognition. In four of the six models, the binding-region nucleotide sequence matched the CTAGCCCCTA motif of the A39 lead aptamer (Fig. 1A, C, D, E). The two exceptions were the truncated (A39T) and a complementary primer containing variant (A39CN) (Fig. 1B, F). The A39CN aptamer lacks the common binding motif present in A39, A39M, A39C, A39CS and, consequently, cannot interact with GbpC through this sequence. Instead, A39CN binds to GbpC via the CTCCCCCTCG motif. Unexpectedly, AF3 predicted that A39T retained its GbpC-binding activity through the same motif identified in A39CN. This observation suggests that the stem-loop structure is responsible for presenting the binding motif in the original A39 aptamer and thus essential for target recognition. Notably, the two distinct binding motifs identified across the six aptamers are both enriched in cytosine residues, highlighting the likely importance of cytosine-rich sequences in mediating GbpC binding.

To further characterise the accuracy of complexes, we analysed the confidence metrics of predicted models provided by AF3 (Table 2.). Each modelling presented moderately high overall accuracy (0.77<pTM<0.85), indicating that the predicted folding of complexes might be similar to the true structure [30,31]. However, the ipTM scores, which measures the predicted relative position of individual subunits within the complex and is therefore a more relevant indicator of multi-component predictions [32], show a greater variability. Derivatives that did not contain the CTAGCCCCTA-motif showed the lowest ipTM values (ipTM(A39T) = 0.65; ipTM(A39CN) = 0.56), whilst models containing the motif yielded moderately high values, most notably the motif itself, which was scored the highest (ipTM(A39M)= 0.93). These suggest that CTAGCCCCTA-motif can create more confidently predicted interfaces with GbpC target molecule, than other segments.

### 3.2. Affinity of A39 aptamer derivatives

Following the in silico predictions, we sought to determine whether the predicted structures indeed form bimolecular complexes. To this end, we employed MST, which enabled the experimental validation of interactions between the A39 aptamer and its derivatives and the GbpC target protein in a homogeneous liquid environment in vitro.

The in vitro characterization of aptamer-protein interactions revealed a crucial role for the primer region, as the K_D_ value of the A39T-GbpC interaction was more than twofold higher than that of the parental A39 aptamer (20.7 ± 2.6 µM vs. 8.09 ± 0.7 µM). A39 derivatives carrying complementary primer sequences, A39C and A39CS, also exhibited clear interactions with GbpC (Fig. 2). However, their affinities were substantially reduced compared with that of the lead aptamer, with K_D_ values of 37.7 ± 4.5 µM and 27.6 ± 2.6 µM, corresponding to 4.6-fold and 3.4-fold decreases in binding affinity, respectively (Table 2). Although the shortened A39 variant lacking part of the random region was not experimentally tested, the stronger binding of A39CS than A39C suggest that the six nucleotides absent from the loop sequence are unlikely to be essential for GbpC binding. In contrast to the AF3 predictions, neither the motif-free derivative A39CN nor the CTAGCCCCTA motif construct A39M exhibited detectable binding to GbpC.

**Figure 2.**
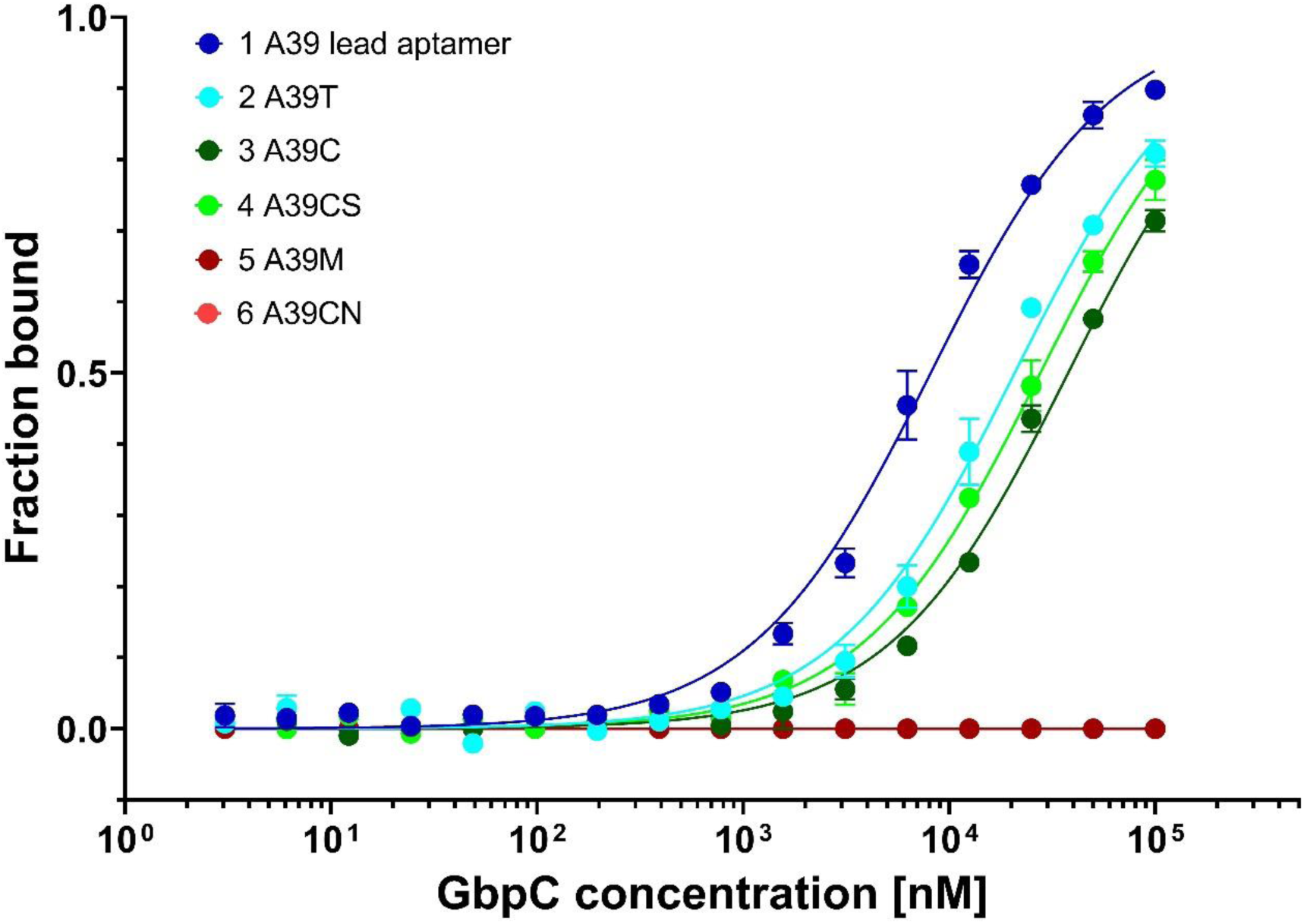
Microscale thermophoresis (MST) data of GbpC-aptamer derivatives. The baseline corrected normalized fluorescence, normalized for amplitude results (fraction bounds) of A39 and its derivatives are plotted. Binding affinities were acquired of GbpC-aptamer interactions by fitting Kd Model of MO.Affinity Analysis software. Data are presented as mean ± standard deviation from at least two individual repeats.

These findings suggest that additional structural features of the aptamer-GbpC complex may contribute to the binding. Alternatively, regions outside the predicted protein-binding motif may be required to establish the three-dimensional architecture necessary for the correct presentation and accessibility of the binding motif.

## 4. Discussion

In this study, the interaction between GbpC and its specific aptamer was analysed using AlphaFold 3 to gain structural insights for the further optimisation of the original A39 aptamer. The crystal structure of streptococcal GbpC has previously been resolved, revealing its experimentally validated calcium-binding residues and a putative glucan-binding pocket [33]. The AF3-predicted complex localised the aptamer-protein interaction within this ligand-binding groove, placing the aptamer in a vicinity of a calcium ion. This finding suggests that both the local electrostatic environment and divalent cation coordination may contribute to aptamer recognition and binding. Notably, the predicted binding site overlaps with the glucan-binding pocket, supporting our previously reported observation that the aptamer and dextran compete for binding to GbpC [25].

To validate the functional importance of the nucleotide motif identified by AF3 and to optimise the lead aptamer, five additional aptamer variants were designed. Interestingly, AF3 localised the binding site of all aptamers to the glucan-binding pocket of GbpC. Moreover, four of the six aptamers shared the same nucleotide motif that was predicted to mediate binding in the original aptamer. In contrast, the A39CN aptamer, which lacks this conserved binding motif, was predicted to interact with the protein through an alternative nucleotide sequence. Surprisingly, the primer-truncated A39T aptamer was also predicted to bind through the very same motif as those of A39CN.

Although revealing the detailed molecular-level spatial structure of aptamer-complexes facilitates optimisation of aptamers, in vitro experimental validation of the putatively improved variant remains indispensable before their further application. To this end, we employed MST to characterise the GbpC-binding properties of the aptamers studied in silico. The measurements demonstrated that none of the modified aptamers outperformed the parental A39 aptamer in terms of binding affinity. Instead, they exhibited dissociation constants that were 2.5- to 4.5-fold higher than that of A39. Furthermore, the two aptamers lacking the predicted glucan-binding pocket-interacting motif present in the other four variants showed no measurable binding to GbpC. These findings suggest that the CTAGCCCCTA motif, predicted by AF3 to interact directly with the protein, is essential for GbpC recognition, whereas additional structural elements of A39 appear to contribute to and strengthen this interaction.

The experimentally obtained results were not fully in concert with the AF3 predictions. The A39CN aptamer, which lacks the binding motif present in the parental aptamer, was necessarily predicted to interact with GbpC through an alternative motif. However, no interaction was detected in the MST measurements. In contrast, the primer-truncated parental aptamer A39T clearly interacted with its target in vitro, yet it was predicted to bind GbpC via the same alternative motif as A39CN. These contradictory observations call into question the validity of the proposed alternative motif as the GbpC-binding region of the aptamer. Notably, the corresponding AF3 models exhibited the lowest ipTM scores among the predicted complexes (0.56 and 0.65, respectively). Conversely, the model with the highest ipTM score corresponded to A39M, the shortest aptamer examined and the only variant retaining exclusively the parental binding motif; however, this aptamer failed to show detectable interaction with GbpC in the MST experiments. These findings indicate that the parental motif is neither strictly sufficient nor entirely dispensable for GbpC binding. Moreover, the lack of agreement between AF3 predictions and the experimental data suggests that neither the predicted binding motifs nor the associated ipTM scores reliably reflected the binding behaviour of these aptamer-protein complexes. By using a broader pool of aptamer derivatives, hitherto undiscovered correlations between predictions and affinity might also be identified, though this would require a significant investment of financial and time resources.

The simultaneous application of structural prediction and interaction analysis methods is becoming increasingly common in molecular biology research. For instance, a direct cryo-EM validation, supported by ITC measurements, demonstrated the importance of experimental confirmation for a ternary protein complex predicted by AF3 [34]. Similarly, the folding mechanisms of protein variants have been successfully assessed using AF3 in combination with additional biophysical methods, such as MST and optical tweezer assays [35]. On the other hand, limitations of AF3 also have been reported, indicating the predicted structures may correlate poorly with experimentally derived thermodynamic parameters [36]. A recent publication evaluated the accuracy of aptamer-target predictions in a comparative study and pointed out that the spatial conformation of aptamers that have not been experimentally solved is still a considerable challenge for AF3 [15]. AF3 performed well when the predicted aptamer-complexes were compared to experimental determined spatial structures but was less reliable with predicting non-canonical nucleic acid secondary structures, which are not represented in the training space. Notwithstanding, the authors suggest that AF3 is feasible to localize the binding sites in aptamer–protein and aptamer–small-molecule complexes [15]. Another systematic evaluation was more critical of deep learning-based structure prediction approaches, reporting that these models frequently place ligands into protein binding pockets despite clearly unfavourable electrostatic or steric interactions. The authors therefore emphasized the need for caution when applying such models in drug discovery and protein engineering [37]. Our findings are consistent with these observations. All aptamers examined in the present study were predicted to bind the glucan-binding region of GbpC, even though two of them showed no detectable interaction with the target in vitro.

In summary, our observations suggest that AlphaFold 3 is a valuable tool for generating hypotheses regarding aptamer-target interactions, but reliable prediction of such interactions remains in its early stages. Consequently, AF3-derived models should be interpreted with caution and their predictions validated experimentally. As the number of experimentally resolved aptamer-target structures continues to grow, the predictive performance of these models is expected to improve. This underscores the importance of expanding structural databases with high-quality aptamer-target complex data to enhance model training.

## 5. Acknowledgements

This work was supported by the 2025–2.1.1-EKÖP-2025–00014 University Research Scholarship Programme of the Ministry for Culture and Innovation from the source of the National Research, Development and Innovation Fund.

This work was funded by the Hungarian National Research, Development and Innovation Office (NKFIH grant No. ANN-139564 and 2024-1.1.2-NAGYVÁLL_FÓKUSZ). Project No. TKP2021-EGA-24 has been implemented with support provided by the Ministry of Innovation and Technology of Hungary from the National Research, Development and Innovation Fund and financed under the TKP2021-EGA funding scheme. We express our gratitude for the support of Richter Gedeon Research Grant Program (RG-IPI-2024-TP18/058) as well.

## Abbreviations

A39: GbpC-aptamer
AF3: AlphaFold 3
ASB: artificial saliva buffer;
cryo-EM: cryo-electron microscopy
GbpC: glucan-binding protein C
GC%: guanine-cytosine percentage
ITC: isothermal titration calorimetry
K_D_: equilibrium dissociation constant
MST: microscale thermophoresis
NMR: nuclear magnetic resonance
pLDDT: predicted local distance difference test
pTM: predicted template modelling score
ipTM: interface predicted template modelling score
T_M_: melting temperature

## 8. CRediT authorship contribution statement

**Ákos Harkai:** Conceptualization, Data curation, Formal analysis, Funding acquisition, Investigation, Methodology, Project administration, Visualization, Writing – original draft, Writing – review & editing

**Tamás Mészáros:** Funding acquisition, Resources, Writing – original draft, Writing – review & editing

